# Evolution transforms pushed waves into pulled waves

**DOI:** 10.1101/266007

**Authors:** Philip Erm, Ben L. Phillips

**Author notes:** corresponding author, School of BioSciences, The University of Melbourne, Victoria, Australia. School of BioSciences, The University of Melbourne, Victoria, Australia (Ben L. Phillips).

**Keywords:** Allee effect, biological invasion, evolution, pushed/pulled wave, invasion speed

## Abstract

Understanding the dynamics of biological invasions is crucial for managing numerous phenomena, from invasive species to tumours. While Allee effects (where individuals in low-density populations suffer lowered fitness) are known to influence both the ecological and evolutionary dynamics of invasions, the possibility that an invader’s susceptibility to the Allee effect might itself evolve on an invasion front has received almost no attention. Since invasion fronts are regions of perpetually low population density, selection should favour vanguard invaders that are resistant to Allee effects. Evolution in response to this pressure could cause invasions to transition from pushed waves, propelled by dispersal from behind the invasion front, to pulled waves, driven by the invasion vanguard. To examine this possibility, we construct an individual-based model in which a trait that governs resistance to the Allee effect is allowed to evolve during an invasion. We find that vanguard invaders rapidly evolve resistance to the Allee effect, causing invasions to accelerate. This also results in invasions transforming from pushed waves into pulled waves, an outcome with consequences for predictions of invasion speed, the population’s genetic structure, and other important behaviours. These findings underscore the importance of accounting or evolution in invasion forecasts, and suggest that evolution has the capacity to fundamentally alter invasion dynamics.

## Introduction

Biological invasions are ubiquitous. Although typically concerned with invasive organisms, the study of biological invasions applies to a much larger range of phenomena, including species responding to climate change (Thomas et al. 2004), the spread of favourable alleles through a population (Fisher 1937; Barton 1979), the spread of pathogens (Perkins 2011), and the growth of tumours (Orlando et al. 2013). Managing these diverse phenomena requires reliable projections of the speeds at which populations are likely to spread (Travis and Park 2004; Gallien et al. 2010); to these ends, as well as for broader understanding, a wealth of ecological and mathematical approaches have been developed for modelling spreading populations (Shigesada and Kawasaki 1997; Lewis et al. 2016). However, while these models can predict the dynamics of real invasions, they are often unreliable in even the simplest of invasive settings (Andow et al. 1990; Williamson 1999; Hastings et al. 2005; Melbourne and Hastings 2009).

Such failures may in part be due to an historically limited appreciation of the importance of evolution in influencing invasion dynamics. A growing body of theory argues that traits governing the dispersal and reproduction of invaders – the two fundamental determinants of an invasion’s speed (Fisher 1937; Kolmogorov et al. 1937; Skellam 1951) – should be under strong selective pressures on invasion fronts. These selective pressures involve both standard natural selection, operating to increase reproductive rates on the *r*-selected invasion front (Phillips 2009), and ‘spatial selection’, likewise operating to increase dispersal rates through the evolutionary effects of spatial sorting (Travis and Dytham 2002; Shine et al. 2011). That spatial selection results in increased dispersal ability has been convincingly supported by empirical evaluations in both natural and laboratory settings (Cwynar and MacDonald 1987; Hughes et al. 2007; Urban et al. 2008; Ochocki and Miller 2017; Weiss-Lehman et al. 2017); that natural selection likewise leads to an increase in the reproduction rates of invaders has been far less clear. Front line invaders in controlled invasions of bean and red flour beetles exhibited a marked increase in dispersal amongst front line invaders, but no increase in fecundity (Ochocki and Miller 2017; Weiss-Lehman et al. 2017). Conversely, in natural invasions, comparisons between source and colonist populations have often shown the latter to possess a range of traits associated with *r*-selection, albeit with notable exceptions (Bossdorf et al. 2005; Amundsen et al. 2012; Hudson et al. 2015). Usually this conspicuous absence is attributed to potential trade-offs between reproductive ability and other traits (Burton et al. 2010), however actually detecting such trade-offs has proven to be difficult (Chuang and Peterson 2016).

The Allee effect, and its probable impact on selection, may in fact be an unrecognised but important mechanism for regulating the evolution of reproduction on invasion fronts. The Allee effect is a phenomenon in which the increasing density of a population positively correlates with the increasing fitness of its members (*i.e*., positive density dependence; Allee 1958; Stephens et al. 1999). It may be caused by any number of constraints typically faced by individuals exposed to low population densities, such as an inability to find mates, increased vulnerability to predation, and inbreeding (Courchamp et al. 2008; Luque et al. 2016). Fitness may become impaired by the presence of either a weak or strong Allee effect, with the former lowering it such that population growth is slowed, and the latter diminishing it to the extent that a population shrinks when below a critical size, called the Allee threshold (Stephens et al. 1999). Since invasion fronts are by nature regions of low population density, Allee effects may curb the reproduction of front line invaders for species that are subject to them. Apart from slowing the invasion, this would impose intense selection on traits that contribute to the emergent Allee effect. Thus, it appears plausible that invasion fronts themselves could select for individuals with increased resistance to Allee effects, and, by extension, increased low-density fecundity.

Despite this insight, the evolutionary responses of invaders to Allee effects remain relatively unexplored, as do the impact of such evolution on invasion dynamics. Modellers have long recognised the non-evolutionary importance of Allee effects to invasions, where it has been shown that low-density invasion fronts coupled with Allee effects may considerably slow or even stop the advance of invasions (Lewis and Kareiva 1993; Kot et al. 1996; Keitt et al. 2001; Taylor and Hastings 2005). There has also been ongoing interest in the influence of Allee effects on the successful establishment of invasive species, which has extended to evolutionary considerations (Drake and Lodge 2006; Kanarek and Webb 2010; Kanarek et al. 2015). It has furthermore been appreciated that Allee effects may alter evolutionary trajectories on invasion fronts by both providing selection against long-distance dispersal and changing effective population sizes (Hallatschek and Nelson 2008; Burton et al. 2010). There has not, however, been a substantial attempt to examine the possibility that Allee effects themselves may evolve, nor how this might influence overall invasion dynamics.

Any evolution of Allee effects would have the potential to reshape the fundamental structure of an invasion. Allee effects are particularly important in that they control the two major classes of invasion: pushed and pulled waves (van Saarloos 2003; Barton and Turelli 2011; Lewis et al. 2016). Pushed invasion waves are primarily driven by individuals from behind the invasion front, and are typical of populations subject to a strong Allee effect where growth is highest at intermediate densities (Lewis et al. 2016). Pulled invasion waves on the other hand are driven by individuals on the leading edges of invasions, where growth can be high if a population is instead subject to a weak or even no Allee effect (Lewis et al. 2016). This relationship between the Allee effect and pushed and pulled waves has been empirically demonstrated by Gandhi et al. (2016), who were able to produce both pushed and pulled yeast invasions by subjecting invaders to different strengths of Allee effect. As Allee effects may be expected to exercise selection on the low-density fecundity of front line invaders, there exists a plausible mechanism through which a single invasion wave could itself transition between wave classes. If front line invaders evolve higher low-density fecundity through increased resistance to the Allee effect, then what began as a pushed wave could potentially transform into a pulled wave, fundamentally altering the dynamics of an invasion as it progresses.

To examine these possibilities, we develop a simulation model of an invasion with heritable variation in individual resistance to an Allee effect. We observe whether invasions can exert selection on a trait that governs the strength of the Allee effect, and whether the resultant evolutionary response is sufficient to have an invasion transition from a pushed to pulled wave.

## Methods

### General description of model

We developed an individual-based model in which invaders were tracked as they dispersed and reproduced across a one-dimensional landscape of patches (Fig. A1). Both time and space were considered discrete. Generations were non-overlapping, with each cohort of invaders dying after one opportunity at reproduction and dispersal. All invaders in the founding generation were randomly assigned a value for a trait (*A*) that determined their Allee threshold, and so reproductive output at different densities. This trait also governed a fitness trade-off across high and low densities, such that individuals adapted to low densities were disadvantaged at high densities, and vice-versa. All individuals reproduced clonally, with offspring receiving their parent’s *A* trait. The trait was, however, allowed to mutate in randomly selected offspring. All simulations were performed using R version 3.5.0 (R Core Team 2018). The model code in its entirety can be accessed at https://github.com/PhilErm/allee-evolution.

### Population dynamics

Reproduction was described by a modified version of a growth function subject to Allee effects first used by Haond et al. (2018). This determined each individual’s expected reproductive output *W* (at location *x*) as

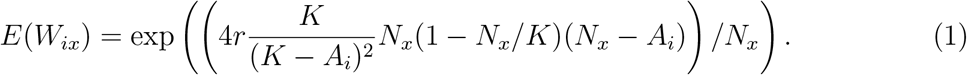

Here, *E*(*W_ix_*) is individual *i*’s expected number of offspring at a particular location *x, r* is the density independent reproductive rate, *N_x_* is the number of individuals at location *x, K* is patch carrying capacity (at densities above which *E*(*W_ix_*) was lower than 1), and *A_i_* the individual’s Allee threshold (at densities below which *E*(*W_ix_*) was lower than 1; Fig 1). As *A_i_* could take any positive or negative value (∈ ℝ), Eq. 1 ensured that, as an individual’s Allee threshold decreased, so too did its high-density performance (Fig. 1).

**Figure 1:**
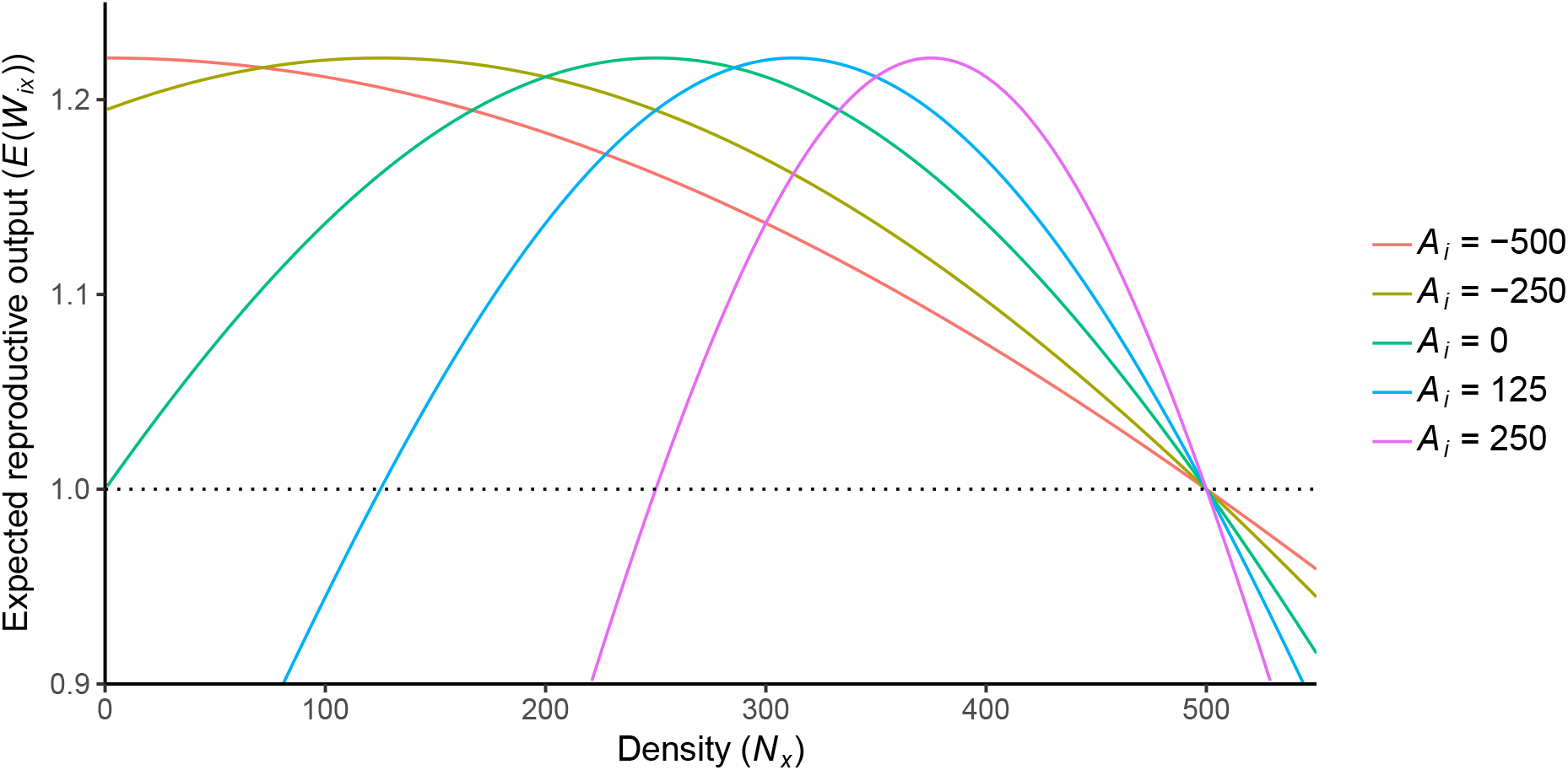
Sensitivity analysis of Eq. 1, a growth function with an Allee effect and reproductive trade-off across densities. *A_i_* determines the Allee threshold, or the critical density below which a population begins to shrink (here seen for values of *E*(*W_ix_*) below the dotted line).

To introduce demographic stochasticity and convert the expected number of offspring to an integer value, an individual’s realised reproductive output *W_ix_* was drawn from a Poisson distribution with λ = *E*(*W_ix_*).

### Spatial dynamics

The invasion space was bounded at *x* = 0, with patches taking values of *x* = 0, 1, 2, …. Individuals dispersed with probability *m* to neighbouring patches in the landscape. Those that dispersed moved to either *x* − 1 or *x* + 1 with equal probability. If an individual attempted to move to *x* = −1, they were returned to *x* = 0.

### Trait variation and inheritance

All founding individuals for each invasion were assigned a value for *A_i_* drawn from a Gaussian distribution with *μ* = *Ā_init_* and *σ* = *σ_A_*. This established the standing trait variance at the beginning of each invasion. Except where mutation occurred, *A_i_* was passed with perfect inheritance to any offspring produced.

In simulations with mutation, newborn offspring were randomly selected with probability *p_mut_* and assigned a new *A_i_* value. Their new mutated value was randomly drawn from a Gaussian distribution with a mean set to their pre-mutation *A_i_*, and a standard deviation of *σ_mut_*.

### Invasion simulations

We used a range of parameterisations to investigate the interaction between evolution and Allee effects, as well as to explore the transition of invasions from pushed to pulled waves (Table 1). Most simulations consisted of the same basic scenario: 300 individuals were distributed evenly across patches 0, 1, and 2, and then underwent 250 generations of dispersal and reproduction. 20 such replicate invasions were conducted for each parameterisation.

**Table 1:** Invasion simulation parameterisations. For the sensitivity analysis, parameters were fixed at reference case values as one parameter was explored across the ranges indicated.

| Parameter | Description | Reference case | Sensitivity analysis | Pushed to pulled | Mixing analysis |
| --- | --- | --- | --- | --- | --- |
| $r$ | Density independent reproduction rate | 0.2 | 0.1, 0.2, 0.3 | 0.2 | 0.2 |
| $K$ | Carrying capacity | 500 | 250, 500, 750 | 500 | 500 |
| $\bar{A}_{init}$ | Mean value of Allee trait for initial invaders | 0 | -50, -25, 0, 25, 50 | 250 | 250 / -497 |
| $\sigma_A$ | Standard deviation of Allee trait for initial invaders | 20 | 20 (0 for exploration of $\bar{A}_{init}$) | 10 | 0 |
| $p_{mut}$ | Allee trait mutation probability | 0.1 | 0, 0.001, 0.01, 0.1 | 0.1 | 0 |
| $\sigma_{mut}$ | Standard deviation of Allee trait mutation | 20 | 20 | 100 | 0 |
| $m$ | Dispersal probability | 0.5 | 0.5 | 0.5 | 0.5 |

#### Reference case

The first parameterisation served as a reference case to provide a general impression of how *A_i_* changed over the breadth and duration of the simulated invasions. At the end of each replicate invasion, we recorded the mean *A_i_* value of invaders in each patch (hereafter called *Ā_x_*) and their density *N_x_*. We also recorded *Ā_x_* over both time and space for a single typical realisation of the model.

#### Sensitivity analysis

We next explored parameter space around this reference case by modelling the impact of changes to *r, K, p_mut_*, and *Ā_init_* on the degree of evolution taking place in the invaders. Each parameter was individually varied across the range of values listed in Table 1, whilst the other parameters were fixed at their reference case values (except for the exploration of *Ā_init_*, which used *σ_A_* = 0 to ensure that the initial invaders possessed exact trait values). At the end of the simulations, we recorded mean trait values for both the core and vanguard of each invasion (*Ā_fin_*). The core mean was calculated across individuals that occupied patches 0 to 4, whereas the vanguard mean was calculated across the 5 farthest occupied patches. We also recorded each invasion’s speed over time to see how evolution and each parameter affected spreading dynamics more broadly.

#### The transition from pushed to pulled waves

To determine if evolution in the vanguard was capable of causing invasion waves to transition from pushed to pulled, we undertook a parameterisation in which all replicate invasion waves started as pushed. We achieved this by setting *Ā_init_* to 250 and *σ_A_* to 10, subjecting all founding invaders to a strong Allee effect. This necessitated increasing the starting population size in each initially occupied patch to 500 (up from the default of 100) to ensure that the populations remained above their Allee thresholds and so prevent extinction in the first few generations. We also increased *σ_mut_* to 100 to ensure that any evolution would happen in a tractable time frame. We then tracked *Ā_van_* (the mean *Ā_x_* of the 5 farthest occupied patches) in invasions for 500 generations. If the mean final value for *Ā_van_* resulted in a monotonically decreasing reproductive output across density for Eq. 1, then the invasions were considered to have become pulled waves (Kolmogorov et al. 1937; Gandhi et al. 2016). For our parameterisation, this critical value (hereafter referred to as *Â*) occurred at *Ā_van_* = −497.

#### Mixing analysis

Finally, to confirm that the waves had transformed from pushed to pulled in practice, we examined the genetic mixing of neutral traits for two sets of invasions where all invaders possessed either *A_i_* = 250 (a strong Allee effect) or *A_i_* = −497 (no Allee effect). No mutations of *A_i_* were allowed to occur. For pushed invasion waves in general, genetic diversity stays high on the wave front as many individuals contribute to the colonisation process, whereas for pulled invasion waves, genetic diversity is rapidly eroded due to repeated instances of the founder effect (Roques et al. 2012). To see if these processes were occurring in our own waves, we assigned all initial invaders a perfectly heritable neutral trait determined by the patch that they started in. Individuals starting in patch 0 were assigned trait 0, individuals starting in patch 1 were assigned trait 1, and so on. To allow for a greater number of traits than would exist for our default number of 3 starting patches, initial invaders were instead spread over 5 patches, with 500 individuals per occupied patch. These invasions were run for 500 generations.

To examine whether diversity in the neutral traits on the wave front was generally preserved (as would be expected in a pushed wave) or lost (as would be expected in a pulled wave), we measured their mean diversity over time in the vanguard 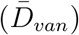 of each invasion using Simpson’s diversity index (Simpson 1949)

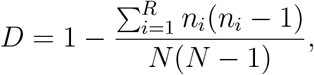

where *D* is the degree of diversity (1 is infinite diversity, 0 is no diversity), *R* is the number of unique traits in the population, *n_i_* is the number of individuals with trait *i*, and *N* is the number of individuals in total. We also recorded the number of individuals carrying each trait in each patch across the whole invasion extent for a single typical realisation of each kind of wave. Since invasions with a strong Allee effect moved much more slowly than those with no Allee effect, we also performed an additional n = 3 simulations of invasions with *A_i_* = 250 that ran for 5000 generations. This meant that they covered the same approximate number of patches as the invasions with no Allee effect that had run for 500 generations.

## Results

### Reference case

*Ā_x_* showed a strong response to selection based on proximity to the invasion front and time since patch colonisation (Fig. 2). Patches on or close to the front exhibited much lower *Ā_x_* values than patches closer to the invasion origin, indicating that increased resistance to the Allee effect had evolved there.

**Figure 2:**
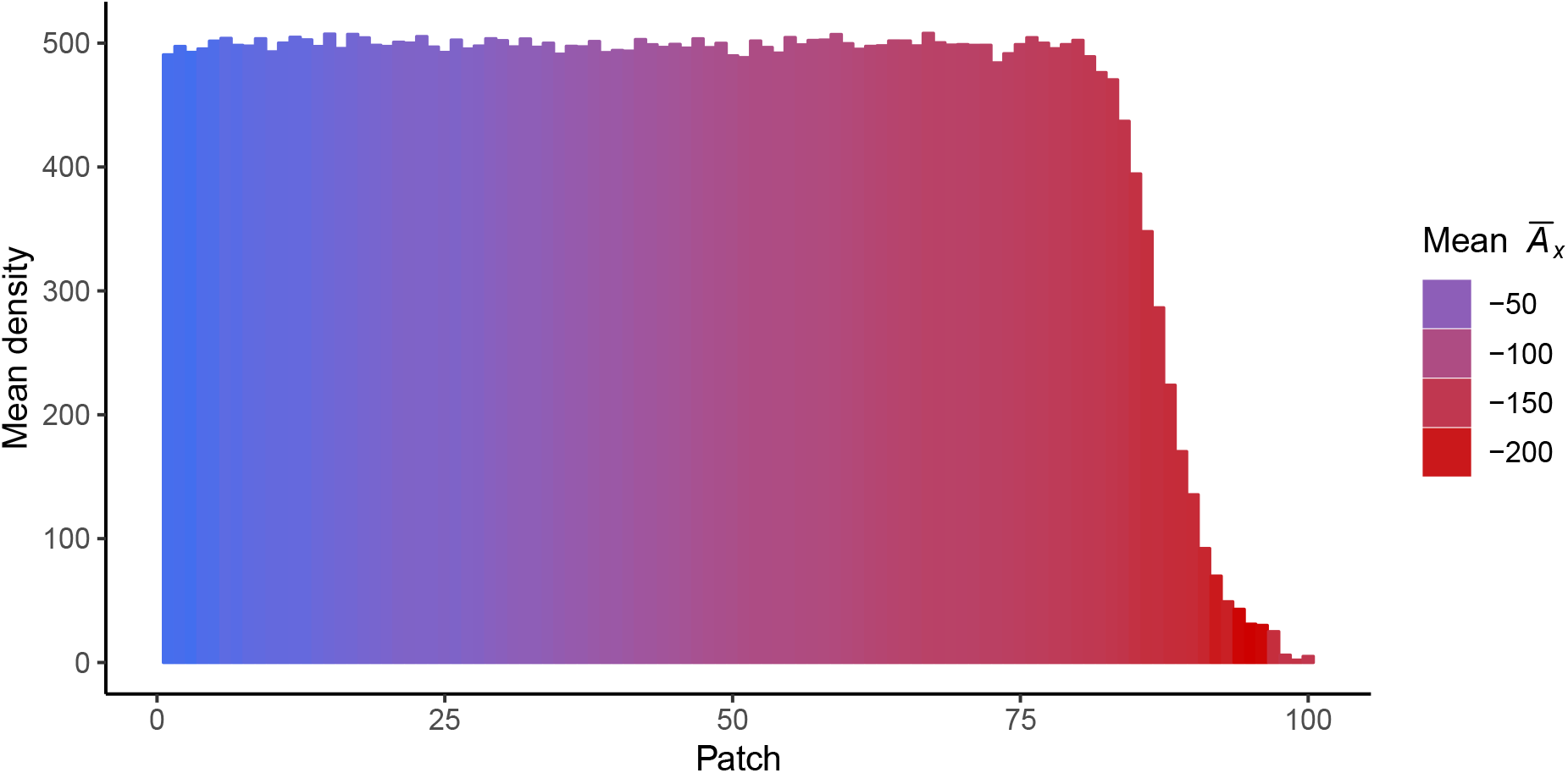
Mean *Ā_x_* (the mean value of *A_i_*, an Allee threshold governing trait, for all individuals located in the same patch) and mean density for invasions (n = 20) after 250 generations under default parameters. More recently colonised patches exhibit greater levels of resistance to the Allee effect, as shown by their lower mean values of *Ā_x_*.

Changes to *Ā_x_* over time in the invasion vanguard took place at a relatively steady rate, but was comparatively inelastic in the invasion core (Fig. A2A). The invasion front in particular was also characterised by highly stochastic deviations in *Ā_x_* driven by instances when the front patch was occupied by just one or a few individuals. There was however no indication of an equilibrium state emerging on the invasion front, with *Ā_x_* still decreasing when simulations ended. Invasions exhibited standard wave profiles over time (Fig. A2B), and, once reaching carrying capacity, patch population densities fluctuated by approximately ±50 individuals around *K*.

### Sensitivity analysis

These basic results appeared to be robust to variation in parameters. Without exception, vanguard individuals always showed a propensity for evolving greater resistance to the Allee effect than did core individuals (Fig. A3). Increases in *r* and *K* accelerated the evolutionary differentiation of the core and vanguard (Figs. A3A and A3B). Increases in *p_mut_* also did so in a more dramatic fashion, with higher mutation rates resulting in much lower *Ā_fin_* values in the vanguard, as well as increased variation in simulation outcomes (Fig. A3C). Even in invasions without mutation, standing variation still enabled differentiation between core and vanguard populations. Changes to *Ā_init_* caused an approximately linear shift in *Ā_fin_* by the end of the simulations (Fig. A3D).

Invasion speeds changed with all parameters (Fig. A4). Increases in *r* led to an increase in speed, whereas the opposite occurred for *K* (Figs. A4A and B). As would be expected if mutation was supplying variance, a high mutation rate resulted in faster and more obviously accelerating invasions, with those using *p_mut_* = 0.1 increasing in speed steadily (Fig. A4C). Decreases in *A_init_* gave rise to invasions that moved faster, but whose ultimate spreading rates nonetheless remained similar to other *A_init_* values (Fig. A4D).

### The transition from pushed to pulled wave dynamics

Invasions readily transformed from pushed to pulled waves over time (Fig. 3). After invasions commenced, *Ā_van_* quickly decreased below 0 (the transition point between a strong Allee effect and a weak Allee effect on the invasion front) and kept decreasing (Fig. 3A). This decrease eventually subsided, with *Ā_van_* stabilising around a quasi-equilibrium value of −497 (*i.e*., *Â*) by the end of the simulations. As *Ā_van_* ≤ *Â* after 500 generations (indicating that Eq. 1 was monotonically decreasing in the vanguard; Fig. 3B), invasions had transformed from pushed waves, subject to strong Allee effects, to pulled waves, subject to no Allee effects at all.

**Figure 3:**
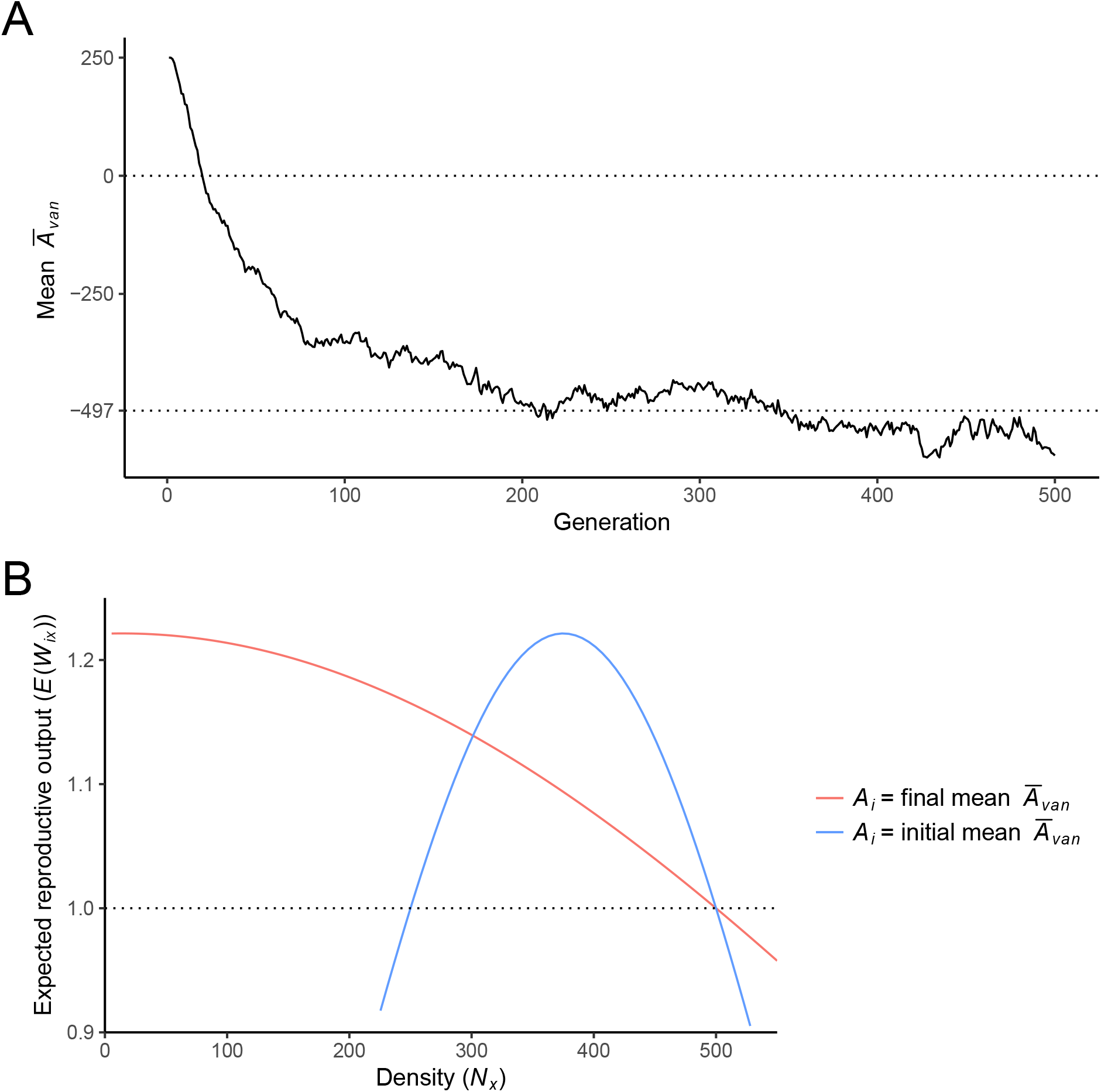
(**A**): The evolution of *Ā_van_* (*Ā* of the vanguard invaders) over time for n = 20 invasions. The founding invaders are subject to a strong Allee effect, meaning that invasions start as pushed waves. The dotted line at mean *Ā_van_* = 0 shows the point at which a transition between a strong (above the line) and weak (below the line) Allee effect occurs. The dotted line at mean *Ā_van_* = 497 shows the point below which no Allee effect occurs. (**B**): A comparison of the expected reproductive output of vanguard invaders at both the beginning and end of the invasions in (**A**). Despite the initial invaders experiencing a strong Allee effect (blue line), by the final generation vanguard invaders have evolved to experience no Allee effect at all (red line). Due to the final generation’s monotonically decreasing reproductive output, pulled invasion waves are theoretically expected to result.

### Mixing analysis

Waves starting as pulled (*Ā_init_* = −497) rapidly lost vanguard diversity whereas those starting as pushed (*Ā_init_* = 250) maintained close to maximum diversity (Fig. 4A). Although both kinds of waves mirrored each other in initially experiencing a sudden loss of diversity, pushed waves quickly recovered and instead experienced a small gradual decrease in 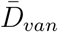 over time. Diversity was still very high 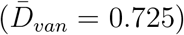 at generation 500, indicating that the pushed waves had maintained a high degree of mixing in the vanguard (Fig. 4B). Pulled waves on the other hand experienced a massive and rapid loss of diversity over time, with all replicates eventually possessing no diversity at all in the vanguard by generation 500 (Fig. 4A). For these waves, one trait eventually dominated the front despite conferring no fitness advantage whatsoever (Fig. 4C).

**Figure 4:**
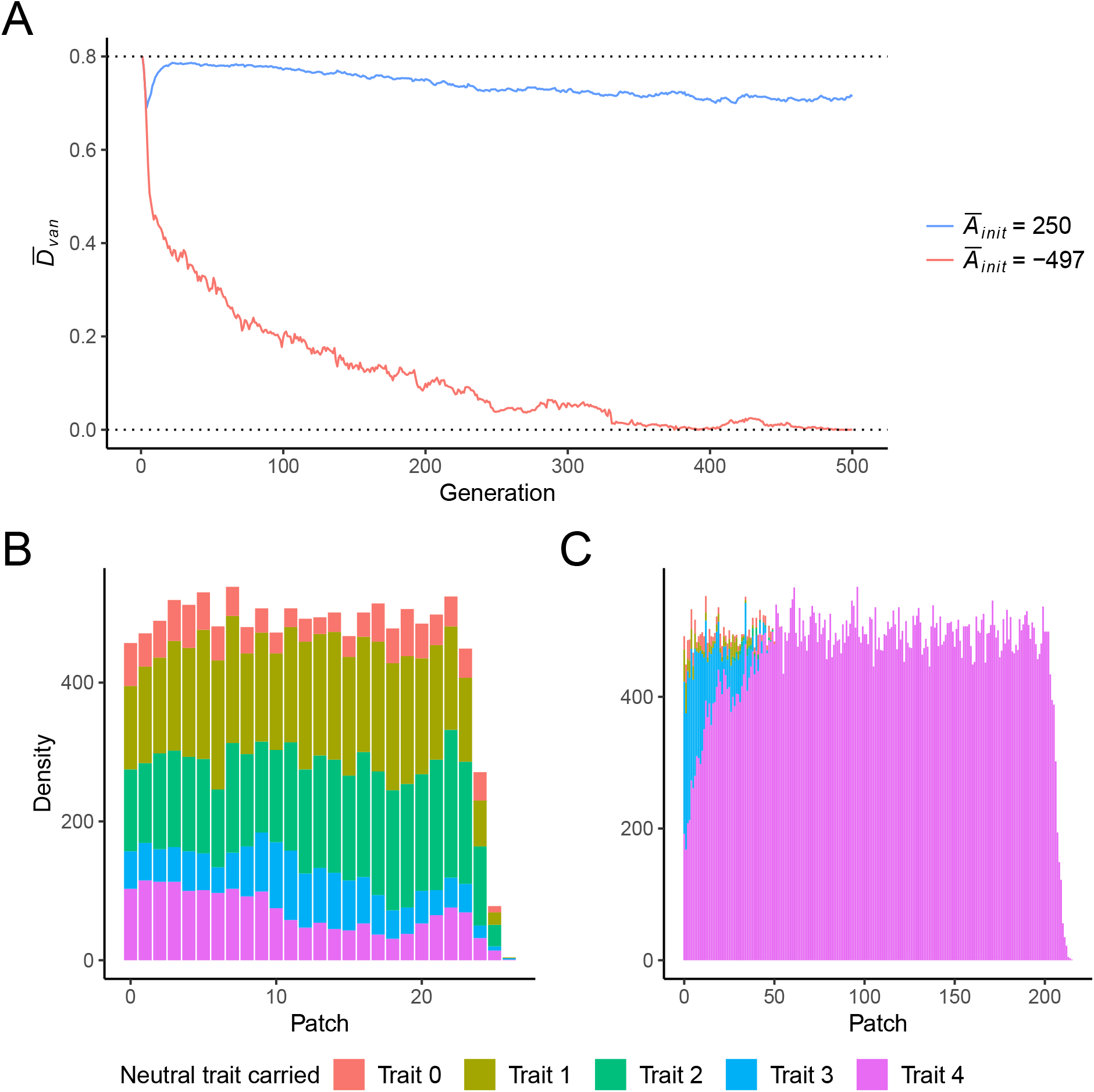
(**A**): The mean neutral trait diversity (as measured by Simpson’s diversity index) over time in the vanguard of n = 20 invasions for two values of *Ā_init_*. Since *A_i_* was not free to evolve, all invaders across all generations possessed *A_i_* values determined by *Ā_init_*. Diversity was maintained in the pushed wave, where a strong Allee effect operated (blue line), but was rapidly lost in the pulled wave, where no Allee effect operated (red line). (**B** and **C**): The structure of neutral trait mixing in the final generation of typical invasion simulations under (*B*) *Ā_init_* = 250 and (**C**) *Ā_init_* = −497. In (**B**), a mix of neutral traits was maintained on the invasion front, whereas in (**C**), despite conferring no fitness advantage, one trait came to dominate.

Pushed waves run for 5000 generations did exhibit a steady decrease in 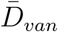 throughout the simulations (Fig. A5A), although even by generation 5000 some diversity was still maintained in the vanguard (Fig. A5).

## Discussion

Due to their importance in dictating the low-density growth rates of stationary and spreading populations, Allee effects have been of long-standing interest to both theoretical and applied ecologists (Boukal and Berec 2002). Here, we consider the possibility that sensitivity to the Allee effect may itself be under selection and evolve in invasions. By permitting invaders to mount an evolutionary response to Allee effects, we show that invasive populations can become resistant to them. As a consequence, invasion waves may transition from pushed to pulled dynamics as Allee effects weaken, with implications for both wave speed and genetic structure.

### Evolved resistance to the Allee effect

In our model, resistance to the Allee effect manifested most strongly on the invasion front, where persistently low population densities appeared to cause ongoing and intense selection for individuals that could reproduce successfully there. Evolution of resistance to Allee effects proved robust to varying initial conditions and key parameters (Fig. A3). Such an outcome is unsurprising; since in all cases the vanguard was propagating into an homogeneous and functionally infinite space, invaders with superior low-density fecundity could easily outcompete rivals. This was in contrast to the invasion core, where resistance remained largely static (Figs. 2 and A3). In these patches, high population densities would have instead ensured that individuals with high Allee thresholds had a competitive advantage. There was however still a slight downward shift in mean *Ā_x_* there; this was likely to have been caused by the particular behaviour of Eq. 1 at densities above *K* (Fig. 1), since individuals with lower *A_i_* values would have momentarily enjoyed increased competitiveness when patch densities oscillated above carrying capacity. Without this peculiarity, it seems likely that resistance to the Allee effect would have decreased in the core, as opposed to remaining mostly static.

Our findings support existing theory about the evolution of life-histories within invasions. The principles of *r*- and *K*-selection, although antiquated in a number of non-trivial aspects, act as useful conceptual tools in this instance (Reznick et al. 2002; Phillips et al. 2010). Since the leading edges of invasions are regions of low population densities, and the long-occupied inner cores of invasions are regions of high population densities, there ought to be a continuum of *r*-to *K*-selective environments from the outer fringes of an invasion back to its point of origin (Phillips et al. 2010). It follows that individuals in the vanguard should evolve traits that enable them to reproduce rapidly at low density, whereas those living in the invasion core should instead evolve traits that enhance competitiveness at high density. In our model, the gradient in *Ā_x_* observed across invasions is strongly concordant with these theoretical expectations (Fig. 2).

This suggests that evolutionary responses to the Allee effect provide yet another life-history axis along which vanguard populations may evolve. The empirical evidence for shifts in reproductive rates in vanguard populations has been mixed (Phillips et al. 2010; Chuang and Peterson 2016; Ochocki and Miller 2017; Weiss-Lehman et al. 2017), but it is entirely possible that these ambiguous results arise from the complex relationship between measurable traits, and actual population growth rates (Reznick et al. 2002). Allee effects introduce additional complexity. In a hypothetical common garden, we might find no difference in seed production, for example, between core and vanguard populations. We might take this as evidence for no shift in reproductive rate. However, it is possible that individuals in the vanguard have more attractive flowers and larger stigmata; a low-density adaptation that, on the invasion front, would see the vanguard individuals exhibit much higher reproductive rates than individuals from the core. We thus need to be very careful about the environment in which we measure reproductive traits, and to be aware that it may be very easy to miss a key trait altogether.

Certainly, direct evidence from invasions themselves appears to support the notion that individuals from recently colonised ranges may have undergone evolution to offset Allee effects. Some organisms that ordinarily exhibit sexual reproduction, such as the parasitic wasp *Mesochorus nigripes*, have instead been found to produce eggs that can hatch even when unfertilised within invading populations (Hung et al. 1988; Hopper and Roush 1993). European starlings (*Sturnus vulgaris*) from recently established colonies have been shown to be more attuned to social signals from conspecifics (Rodriguez et al. 2010), and cane toads (*Rhinella marina*) from the invasion front in northern Australia appear to exhibit increased sociality (Gruber et al. 2017). In both cases, these behaviours enhance each species’ capacity to aggregate despite low population densities. It has been noted that selfing in plants is particularly prominent in marginal populations (Pannell 2015), and it has long been argued that biogeographical biases in the global distribution of parthenogenetic species may in fact reflect their inherent superiority as colonisers (Kearney 2005). Given the relatively straightforward nature of our model’s predictions, further comparisons between core and vanguard populations in nature may prove useful in evaluating its findings.

### Wave type transitions

In the vanguard, the evolution of resistance to the Allee effect was able to progress to such an extent that invasions transitioned from pushed to pulled waves (Fig. 3), a result supported by the mixing analysis in which waves subject to strong Allee effects maintained diversity in their vanguards whereas those subject to no Allee effects did not (Fig. 4). Were this to occur in a real invasion, it would have far-reaching and profound impacts on a population’s structure and dynamics. Because an absence of Allee effects allows populations to grow from small sizes, demographic stochasticity on the invasion front becomes a more powerful force as Allee effects diminish. This stochasticity not only results in intrinsically more variable invasion speeds (Taylor and Hastings 2005; Melbourne and Hastings 2009), but also has several evolutionary consequences. The increased demographic stochasticity and success of small founder populations mean that pulled invasions experience substantial genetic drift – a force that should be strongly mitigated in pushed invasions (Slatkin and Excoffier 2012). In pulled invasions, this powerful drift can cause ‘gene surfing’, where deleterious alleles persist in the vanguard and are spread over wide geographical areas (Hallatschek and Nelson 2008; Graciá et al. 2013), ultimately contributing to populations suffering from high levels of ‘expansion load’ (Peischl et al. 2015). This expansion load can also affect the very traits responsible for spread rate – dispersal and reproduction – such that evolutionary stochasticity makes a very large contribution to making invasion speeds unpredictable (Phillips 2015; Ochocki and Miller 2017; Weiss-Lehman et al. 2017). All of these effects are mitigated on pushed invasion fronts. Here the requirement for large founding populations maintains relatively high levels of genetic diversity (Hallatschek and Nelson 2008; Roques et al. 2012), an expectation reflected in Fig. 4. If invasions become more pulled with time, then all of the stochastic outcomes attendant on pulled waves will increasingly manifest as invasions progress.

Despite the above however, even pushed waves saw a gradual loss of diversity in both the vanguard and the entire breadth of the population if simulated for 5000 generations (Fig. A5). In the present instance this is unsurprising. Although pushed waves driven by strong Allee effects may be expected to maintain diversity indefinitely in an idealised scenario free of genetic drift (Roques et al. 2012), in instances like our own, in which stochasticity allows genetic drift to manifest, it should instead be expected that diversity will decrease over time regardless of circumstances (Hallatschek and Nelson 2008). Thus even a pushed wave will experience a loss in diversity, although the difference in diversity loss between between pulled and pushed waves that have covered approximately the same distance (*i.e*., Figs. 4C and A5B, with the latter requiring an extra 4500 generations to do so) is striking.

The evolution of higher reproduction rates as Allee effects diminish would also facilitate the evolution of increased dispersal ability (Perkins et al. 2013). As Allee effects weaken, highly dispersive individuals accumulating on the invasion front through spatial sorting also accrue a fitness benefit, with spatial sorting and natural selection conspiring to drive increasing dispersal rates. Again, as invasions transition from pushed to pulled, we should see evolutionary processes accelerate.

Accounting for the possibility that invasions may transition from being pushed waves to pulled waves would considerably complicate attempts to predict invasion speed, especially given that the equations conventionally used to model the velocities of pushed and pulled waves are different from one another (Lewis et al. 2016). Furthermore, since pushed and pulled waves each favour the deployment of different control techniques (Gandhi et al. 2016), the most effective strategy for managing a particular invasion may also change as the invaders evolve. Since many control techniques exploit the Allee effect (Tobin et al. 2011), accounting for the resultant selection these impose may be prudent. If control techniques strengthen selection for individuals that are resistant to Allee effects, then they may inadvertently contribute to a less controllable and less predictable invasion.

## Conclusion

Our model represents a first step in describing the higher order effects of the evolution of resistance to Allee effects on invasion fronts. Although our results support the hypothesis that invasion fronts ought to select for individuals that are resistant to Allee effects, further theoretical and empirical work is required. In particular, models that incorporate recombination may see slower or stalled evolution depending on the strength of initial Allee effects. Experimental invasions and observational studies of real invasions may also help determine if transitions from pushed to pulled waves can happen on ecologically relevant timescales. Irrespective, future empirical efforts should pay careful attention to the context in which reproductive rate is assessed. If our findings hold in reality, rapid evolution can act to release the handbrake applied by Allee effects, with faster invasions and more dispersive invaders the result.

## Acknowledgements

We thank Lionel Roques and Diana Fusco for their advice on using trait diversity to diagnose pushed and pulled waves. We would also thank two anonymous reviewers for their comments on an earlier draft of the paper.

