## Supplementary material for "Evolution transforms pushed waves into pulled waves"

March, 2019

***, School of BioSciences, The University of Melbourne, Victoria, Australia.
†**, School of BioSciences, The University of Melbourne, Victoria, Australia

### Figures

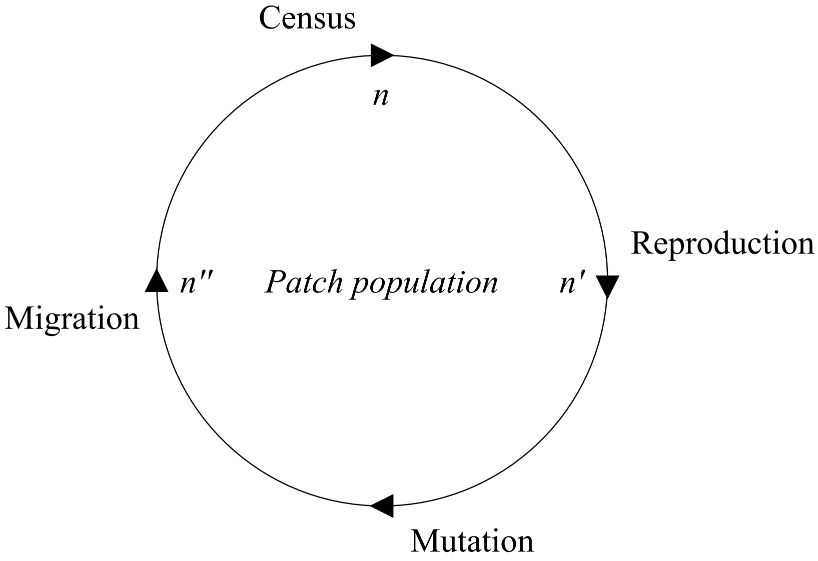


**Figure A1:** The cycle of processes controlling the population size ($n$) of a single patch over one generation. **Census**: The number of individuals in each patch is tallied, and mean trait values recorded. **Reproduction**: Each individual reproduces according to its $A_i$ trait and the density of the patch; all but the newly born offspring perish. **Mutation**: $A_i$ mutates in a random selection of offspring. **Migration**: Offspring randomly disperse to and from the patch. After each generation the population is censused again, and the cycle begins anew.

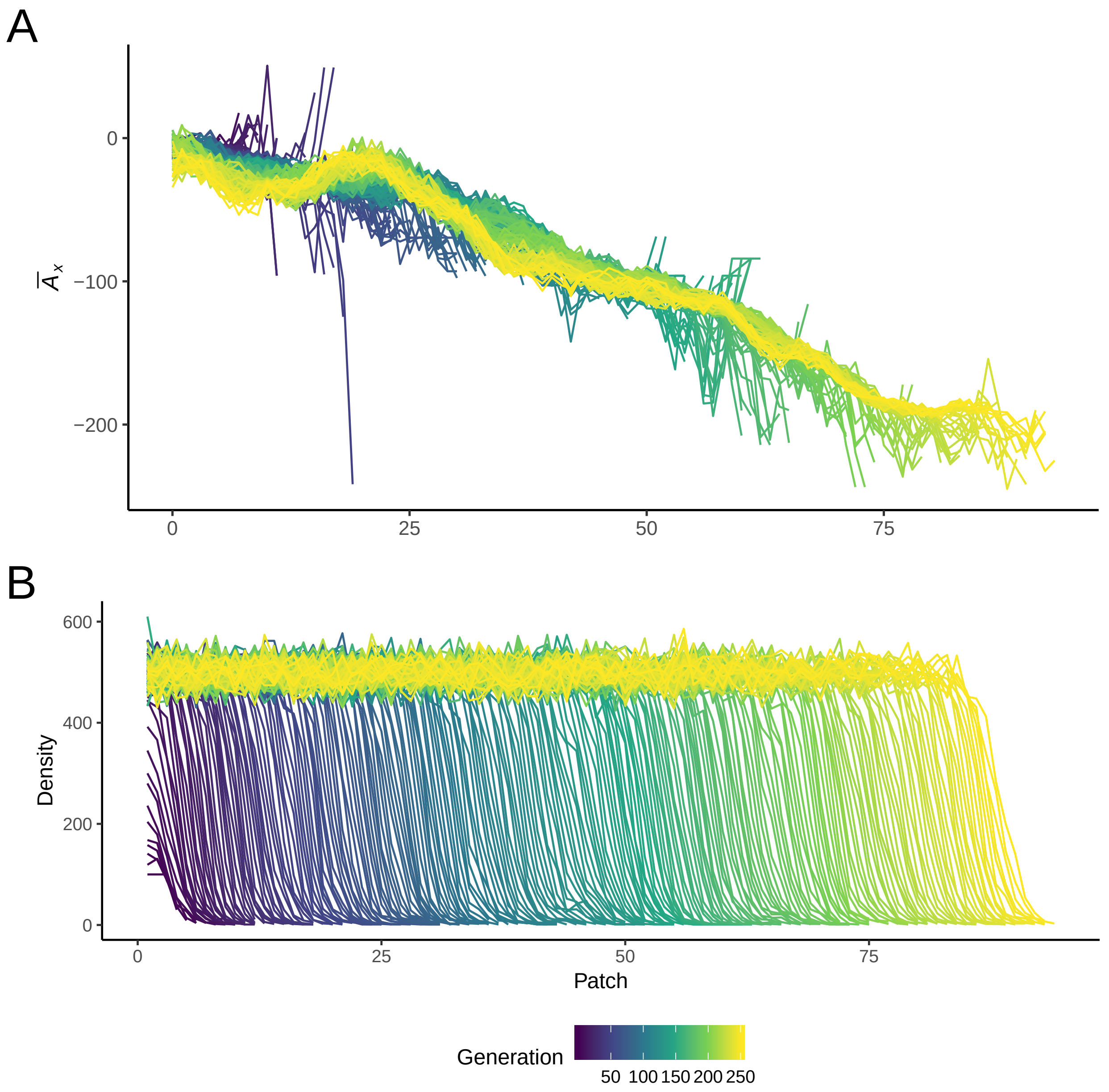


**Figure A2:** (**A**): $\bar{A}_x$ over time and space for a typical invasion under default parameters. Selection on trait variation in the founding generation of invaders and variation introduced over time by mutation results in the differentiation of $\bar{A}_x$ between the core and frontier invaders. (**B**): Density over time and space for a typical invasion under default parameters.

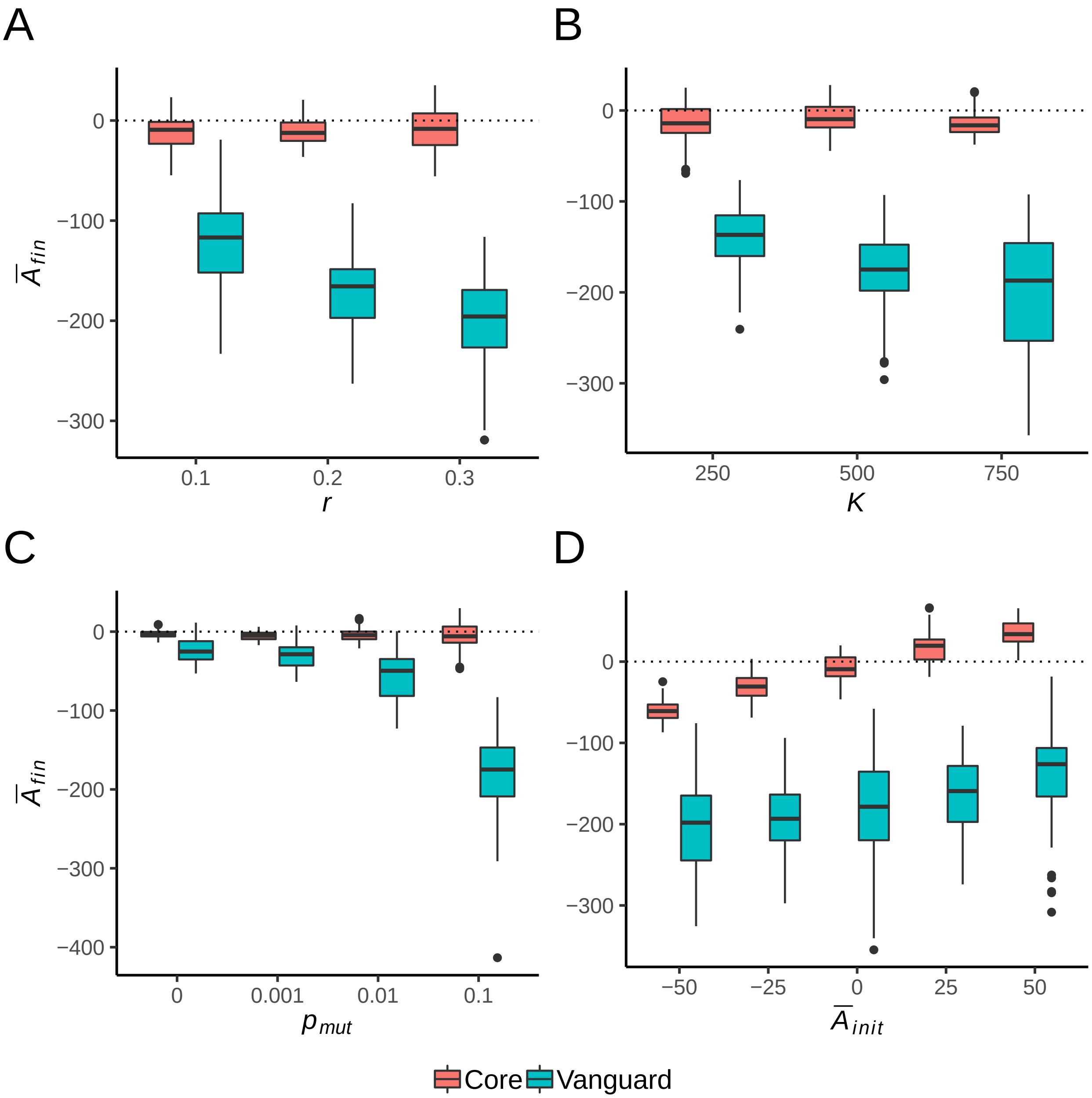


**Figure A3:** $\bar{A}_{fin}$ ($\bar{A}_x$ at the end of an invasion) across parameter space for core and vanguard invaders (n = 20 for each level of each parameter). Core invaders were those occurring in patches 0-4, and vanguard invaders were those within the 5 farthest occupied patches. (**A**) examines reproductive rates, (**B**) carrying capacities, (**C**) probabilities of mutation, and (**D**) $\bar{A}_x$ in the founding generation of invaders. Each parameter was varied while the others were fixed at default values listed in Table 1.

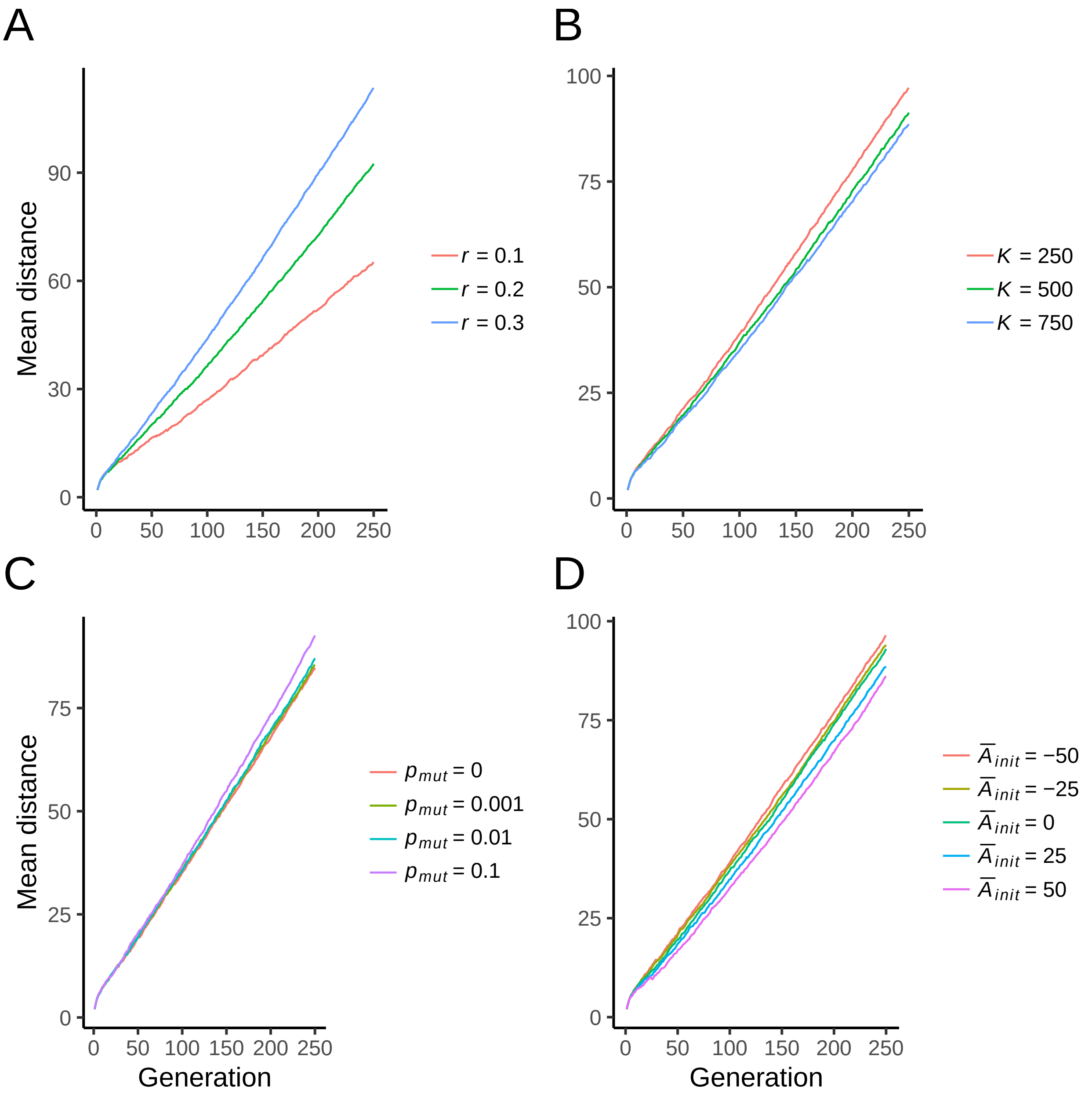


**Figure A4:** The mean distance travelled by invasions (n = 20 per parameter level) across different (**A**) reproductive rates, (**B**) carrying capacities, (**C**) probabilities of mutation, and (**D**) $\bar{A}_{init}$ values. Upwards curving lines denote acceleration.

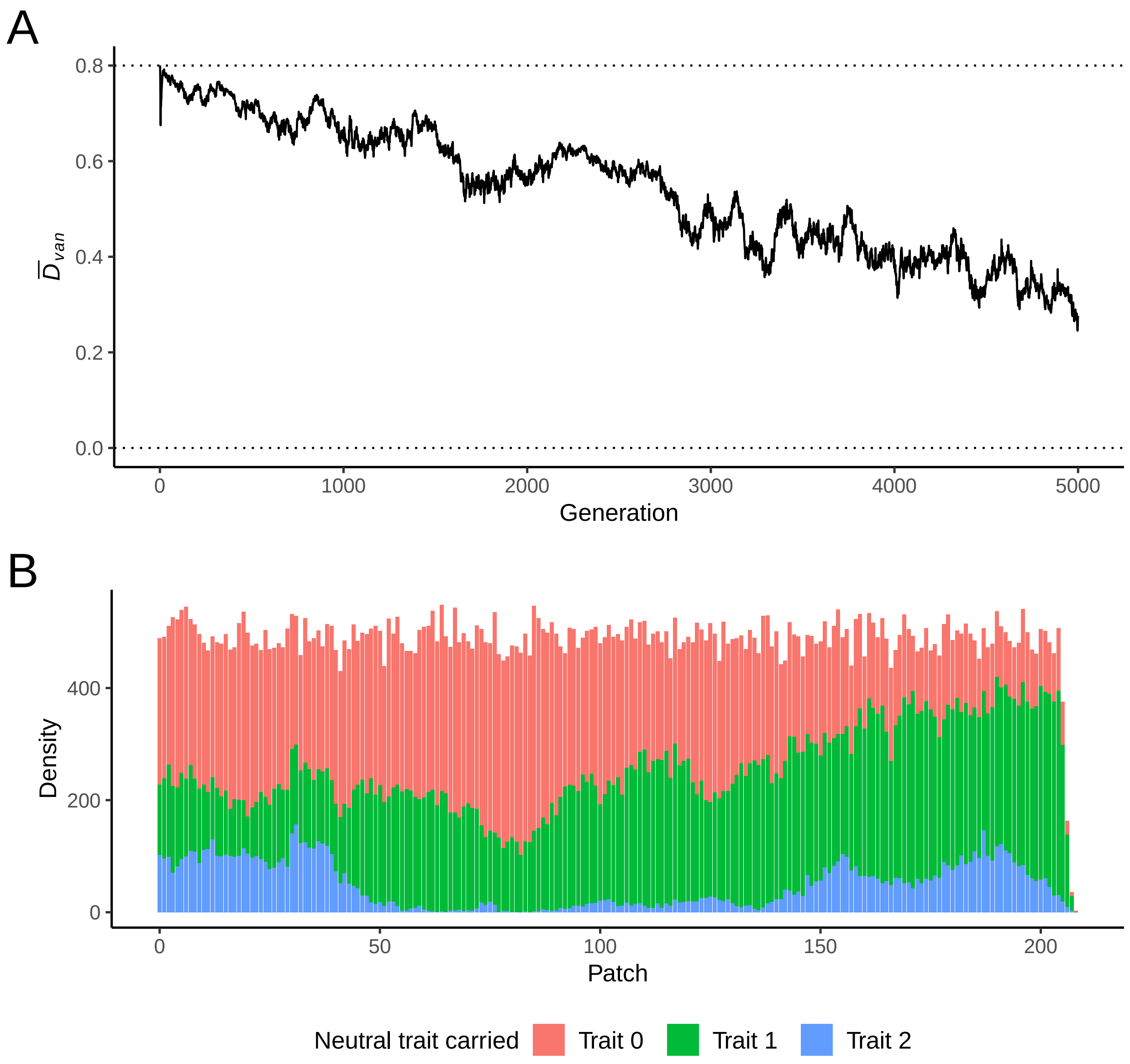


**Figure A5:** (**A**): The mean neutral trait diversity (as measured by Simpson's diversity index) over 5000 generations in the vanguard of n = 3 invasions for $\bar{A}_{init} = 250$. Diversity gradually decreased even under a strong Allee effect. (**B**): The structure of neutral trait mixing in generation 5000 of a typical $\bar{A}_{init} = 250$ invasion simulation. Some diversity is still maintained in the vanguard after 5000 generations, although two traits have been lost to the population entirely. Other particular realisations saw all traits remaining, along with variable numbers of traits on the vanguard at the simulation's end.
